# Single and multi-analyte deep learning-based analysis framework for class prediction in biological images

**DOI:** 10.1101/2022.10.13.512074

**Authors:** Neeraja M Krishnan, Saroj Kumar, Ujjwal Kumar, Binay Panda

**Affiliations:** School of Biotechnology, Jawaharlal Nehru University, New Delhi 110067, India; Special Centre for Systems Medicine, Jawaharlal Nehru University, New Delhi 110067, India

## Abstract

Measurement of biological analytes, characterizing flavor in fruits, is a cumbersome, expensive and time-consuming process. Fruits with higher concentration of analytes have greater commercial or nutritional values. Here, we tested a deep learning-based framework with fruit images to predict the class (sweet or sour and high or low) of analytes using images from two types of trees in a single and multi-analyte mode. We used fruit images from kinnow (*n* = 3,451), an edible hybrid mandarin and neem (*n* = 1,045), a tree with agrochemical and pharmaceutical properties. We measured sweetness in kinnows and five secondary metabolites in neem fruits (azadirachtin or A, deacetyl-salannin or D, salannin or S, nimbin or N and nimbolide or E) using a refractometer and high-performance liquid chromatography, respectively. We trained the models for 300 epochs, before and after hyper-parameters’ evolution, using 300 generations with 50 epochs/generation, estimated the best models and evaluated their performance on 10% of independent images. The validation F1score and test accuracies were 0.79 and 0.77, and 82.55% and 60.8%, respectively for kinnow and neem A analyte. A multi-analyte model enhanced the neem A model’s prediction to ‘high’ class when the D:N:S’s combined class predictions were high:low:high and to ‘low’ class when D:N’s combined class predictions were low:high respectively. The test accuracy increased further to ~70% with a 10-fold cross-validation error of 0.257 across ten randomly split train:validation:test sets proving the potential of a multi-analyte model to enhance the prediction accuracy, especially when the numbers of images are limiting.

## Introduction

Measuring the concentration of an analyte, such as metabolite, enzyme, protein, or any other chemical moiety in biological parts, such as fruits, is a common practice in biology. Each of the analytical (chemical, biochemical, immunological, or imaging-based) methods to measure metabolites provides a different type of readout. Some techniques, although simple to measure, are not precise and others, while accurate and precise, require extensive sample handling, preparation time, expensive reagents and equipment, and specialized skills. Knowing whether the fruit will be sweet or sour may be of practical value to consumers while picking fruits in a supermarket. In other cases, a quick and rough estimate of the concentration class (high or low) of a fruit metabolite may help the industry to choose the right fruit lot to extract chemicals for agrochemical or pharmaceutical use. A quick, inexpensive and easy-to-use method to choose sweeter fruits in the supermarket aisle or fruits with high concentration of a specific metabolite, however ideal, is currently not available.

Color, texture and overall appearance are factors that consumers consider before buying fruits in supermarkets. These are good surrogates for quality, sweetness, and flavor (Clydesdale 1993; Francis 1995). Multiple factors influence these qualities, mainly genetics, climate and environment, which the consumers have no control over, therefore are of little value to the consumers. Therefore, it is desirable to develop an easy-to-use tool to help consumers select fruits that are sweet and flavorful. Additionally, fruit segregation based on their color, taste, and nutrition increases their pre-sales value tremendously. Metabolites, primary and secondary, are intermediate products of metabolic reactions catalyzed by enzymes. Examples of some primary metabolites are amino acids, vitamins, organic acids, and nucleotides. Living cells synthesize most of the primary metabolites essential for their growth. Secondary metabolites, especially those derived from microbial and plant sources, such as drugs, fragrances, dyes, pigments, pesticides and food additives, are not required for the cell’s primary metabolic processes but have a wide range of application in agriculture and pharmaceutical industry. The commercial importance of secondary metabolites has invoked the exploration of producing bioactive plant metabolites synthetically in the laboratory, either by total synthesis, plant tissue culture or metabolic engineering (Keasling 2014). Plant-based metabolites offer a cheaper, non-toxic and environment-friendly alternative to chemical pesticides.

Machine learning and artificial intelligence-based methods have recently found their utility in biology (Minervini et al. 2014, Rostam et al. 2017). Deep learning methods outperform conventional methods of image classification, such as principal components identification, and their usage in Support Vector Machines and Random Forest classifiers (Kovaleva et al. 2002). They are, therefore, more suitable for image-based analyses. The color of the fruit, the only external factor visible to the open eye, provides a unique feature that can be used in computer vision and image analysis. Hence, deep learning-based image classification tools may best predict individual fruit’s overall flavor (due to sugar, acidity and other volatile metabolites).

Here, we have used fruits from two completely different types of trees, kinnow, and neem, to test whether deep learning-based methods can be used to predict the class (sweet or sour in kinnow, and high or low in neem) of metabolites that give the fruit its characteristic taste or value. Kinnow and neem fruits are very different in size, texture, smoothness, color, and shape. Kinnow is a hybrid of two citrus cultivars - King (*Citrus nobilis*) × Willow Leaf (*Citrus deliciosa*), high yielding and has higher juice content than other varieties of oranges. Rich in vitamins (B and C), β-carotene, calcium, phosphorus, kinnows nutritional content are preferred for beverages preparation (Sogi and Singh 2001). Kinnow fruits are preferred over oranges as they are juicier, inexpensive and locally grown in certain parts of south Asia. Image analysis was previously used to identify and assess the quality of fruits, grains, and vegetables (Al-Sammarraie et al. 2022; Boniecki et al. 2015; Zaborowicz et al. 2017; Koszela et al. 2015). In the second case, we used fruits from neem (*Azadirachta indica*) tree. Neem fruits are a source of a wide variety of secondary metabolites, including the potent antifeedant azadirachtin that is widely used as an alternative to chemical pesticide (Schmutterer and Sharma 1997). Neem fruit metabolite concentrations are measured after extraction, biochemistry, and analyses, requiring time, access to expensive equipment, reagents, and specialized skills. Additionally, metabolite concentration often varies widely among plants, even within a small area. Other than the geography of the tree locations, azadirachtin content also depends on the quality of the seeds, age of the trees, soil conditions and climatic factors (Chary 2011; Kaushik et al. 2004; Sidhu et al. 2003; Singh et al. 1999; Priyanka et al. 2010), thus causing the commercially available seeds to yield varying levels of azadirachtin. A previous study has reported high variation, as high as 6-fold, in azadirachtin-A and azadirachtin-B content across seed kernels (Kaushik et al. 2007). To circumvent this, total organic synthesis of azadirachtin was attempted in the past (Veitch et al. 2007). However, the process was time-consuming and produced a low yield (Veitch et al. 2008). Therefore, a quick method to estimate the metabolite concentration class (high or low) will significantly help in selecting trees yielding high metabolite concentrations for the metabolite production process. In addition to azadirachtin (A), we also biochemically measured four other secondary metabolites (deacetyl-salannin or D, salannin or S, nimbin or N and nimbolide or E) from fruits to test whether a multi-analyte deep learning-based object detection-cum-classification framework can boost the accuracy of the model to predict the class of azadirachtin (A).

## Methods

We created deep learning-based single- and multi-analyte frameworks for kinnow and neem fruit images to generate optimum models that can predict analyte classes. We made two classes each for kinnow sweetness W (sweet or sour), and five neem intracellular metabolite concentrations (high or low) for azadirachtin (A), deacetyl-salannin (D), nimbolide (E), nimbin (N) and salannin (S). While the W class prediction was treated as an object detection-cum-classification problem, we added a combinatorial method for predicting A class in neem fruits using the predictions from the D, E, N, and S models to boost the accuracy since all five analytes were measured from the same fruits. The flow of the single- and multi-analyte deep learning framework is outlined in **Figure 1**.

**Figure 1:**
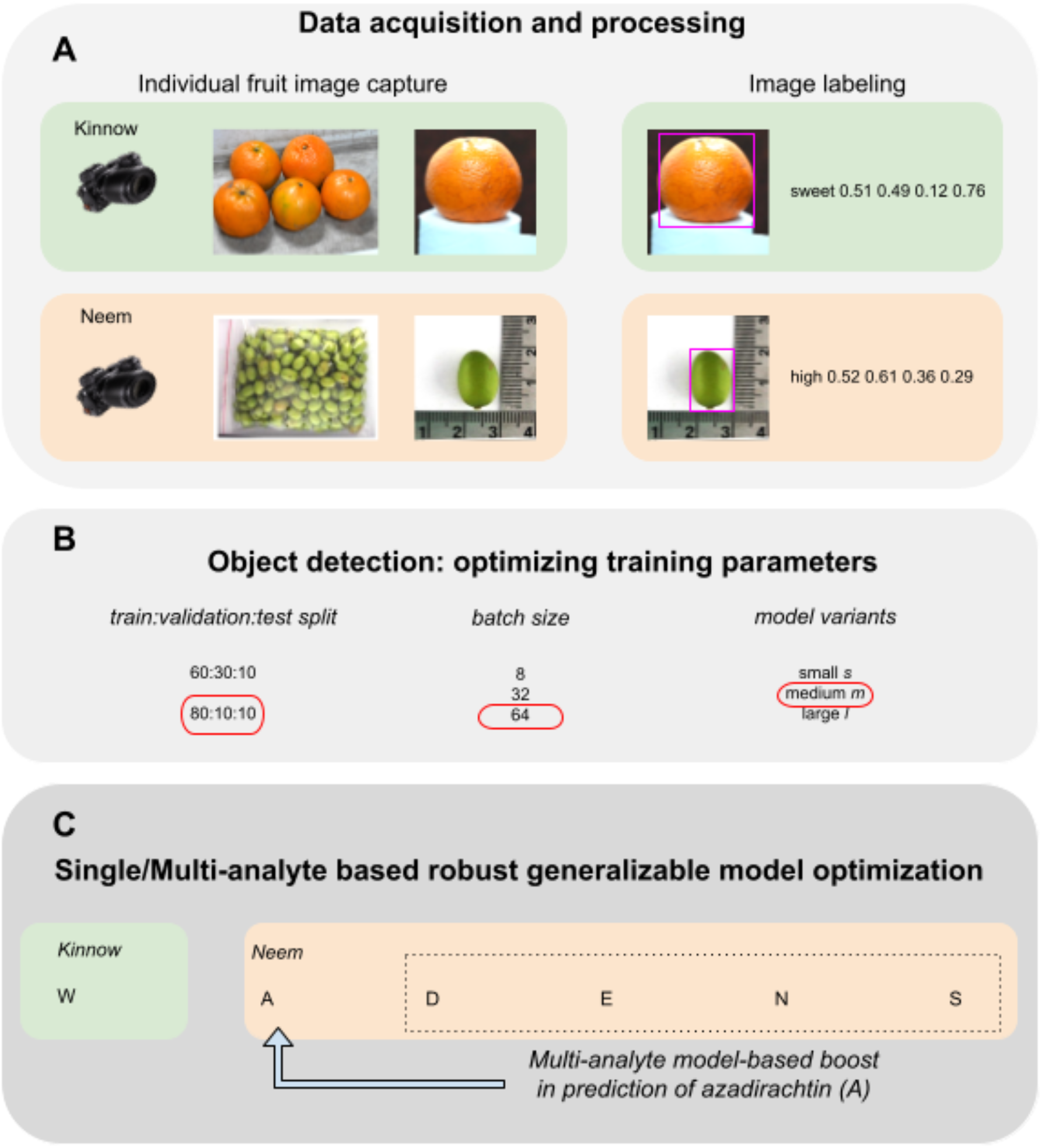
Overview of all the phases of the single- and multi-analyte deep learning framework for class prediction in fruit images.

### Preparation of images, labels and background images

Each fruit was cleaned with water to remove any surface contaminants and wiped dry before image capture. We captured the kinnow fruit images in four orientations per fruit, two side, top and bottom views with controlled backgrounds. We placed the neem fruits sideways and captured the images against white background from a fixed distance. The sweetness level of each kinnow was measured using a handheld digital refractometer and the average of three independent readings was considered. In all, we collected 3,541 kinnow images and labels, with corresponding reading measurements (**Table 1**). For neem fruits, concentrations of A, D, E, N and S metabolites were measured using reverse-phase columns on the high-performance liquid chromatography (HPLC) instrument, with respective analytical standards. The images were divided into ‘high’ and ‘low’ classes for each metabolite using the respective mean concentrations as the thresholds. We drew the bounding boxes around the fruit contours using Makesense.ai (https://www.makesense.ai/) to get the object annotation coordinates. The training images were augmented with ~10% background images chosen randomly from openly accessible and other images (128 images from coco-128; https://www.kaggle.com/datasets/ultralytics/coco128), 54 flower, 50 leaf and 50 fruit images. We also performed imputation experiments using the kinnow dataset by serial and class-wise down-sizing of the dataset by 10%, 20%, 30%,40%, 50%, 60%, 70%, 80% and 90%, performed training with these datasets and studied the trend for the various metrics.

**Table 1:** Number of images within each class for training, validation and test datasets.

| *Fruit (analyte)* | *Class* | *Training* | *Validation* | *Test* |
| --- | --- | --- | --- | --- |
| *Kinnow sweetness (W)* | *Sweet* | 1464 | 1001 | 997 |
|  | *Sour* | 1365 | 965 | 969 |
| *Neem (A)* | *High* | 360 | 50 | 45 |
|  | *Low* | 465 | 75 | 50 |
| *Neem (D)* | *High* | 420 | 45 | 55 |
|  | *Low* | 405 | 50 | 70 |
| *Neem (E)* | *High* | 410 | 55 | 75 |
|  | *Low* | 415 | 40 | 50 |
| *Neem (N)* | *High* | 440 | 40 | 45 |
|  | *Low* | 385 | 55 | 80 |
| *Neem (S)* | *High* | 415 | 50 | 55 |
|  | *Low* | 410 | 45 | 70 |

### Optimization of training parameters and model prediction

We used the YOLOv5 framework (Jocher et al. 2022) for object detection, for training and prediction, due to its high runtime speeds without loss of accuracy. The bounding box labels were exported in the five coordinate YOLOv5 format. Since the neem fruit image numbers were about threefold less in number compared to the kinnow image numbers, we attempted optimization of training parameters using the kinnow data only. We tested combinations of parameters for training such as the split ratio of the dataset into train, validation and test subsets (60:30:10 and 80:10:10), batch size (8, 32 and 64) and also the model itself (small, medium and large) differentiated by the depth of the neural network with 7.2, 21.2 and 46.5 million parameters, respectively. We monitored and recorded the performance metrics, Precision (*P*), Recall (*R*), mAP:0.5, mAP:0.5:0.95 and F1-score 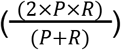 and based the selection of training parameters on maximizing the F1-score. We also monitored the loss curves (object, box and class) for training and validation to prevent over-fitting. We used Weights & Biases (http://wandb.com/; Biewald, 2020) integration with YOLOv5 to track, log and visualize all the runs. We trained all the runs for 300 epochs with the default set of hyper-parameters. Additionally, we performed fine-tuning of hyper-parameters using the ‘evolve’ function for 300 generations and 50 epochs per generation and trained the runs with the evolved hyper-parameters for 300 epochs. After identifying the epoch at which overfitting occurs based on the validation loss curve crossing over the training one, we retrained until that epoch to obtain the best models.

### Model testing

We used the test set of blind images to predict the class for each incoming image and assessed its classification based on the best models in the single-analyte framework for class prediction. For neem fruits, we further identified patterns using the predictions from the D, E, N and S models. We used these patterns to correct the predictions from the A model, and boost its classification accuracy. We termed this as the ‘multi-analyte’ framework for class prediction.

### Cross-validation

We performed 10-fold cross validation on the neem image dataset, as it comprised ~3-fold fewer images when compared to the kinnows. We created ten datasets by performing 80:10:10 random splits of the data into training:validation:testing subsets, ten times. For each of the shuffled datasets, we performed training before and after tuning of hyper-parameters, and tested the prediction efficiency of the best models using the test set images. By comparing the predicted class to the actual class, we obtained the cross-validation error in prediction across the ten shuffled test datasets.

## Results

### Sugar/Analyte distribution

For kinnows, the refractometer readings ranged from 4.87 to 16.53 (**Figure 2A**) with the mean reading ~ 10. Therefore, we labeled kinnows with refractometer readings > 10 as ‘sweet’ and <=10 as ‘sour’. For neem, the concentrations (units) ranged from 0.18 to 1.003, 0.007 to 0.691, 0.056 to 1.420, 0.009 to 0.501, and 0.04 to 0.252, for A, D, S, N and E, respectively, with 0.58, 0.08, 0.145, 0.438 and 0.034, as respective means (**Figure 2B**). We used these mean values as thresholds for concentration values below which the fruits are labeled ‘low’ and above which they are labeled ‘high’, for the respective metabolites.

**Figure 2:**
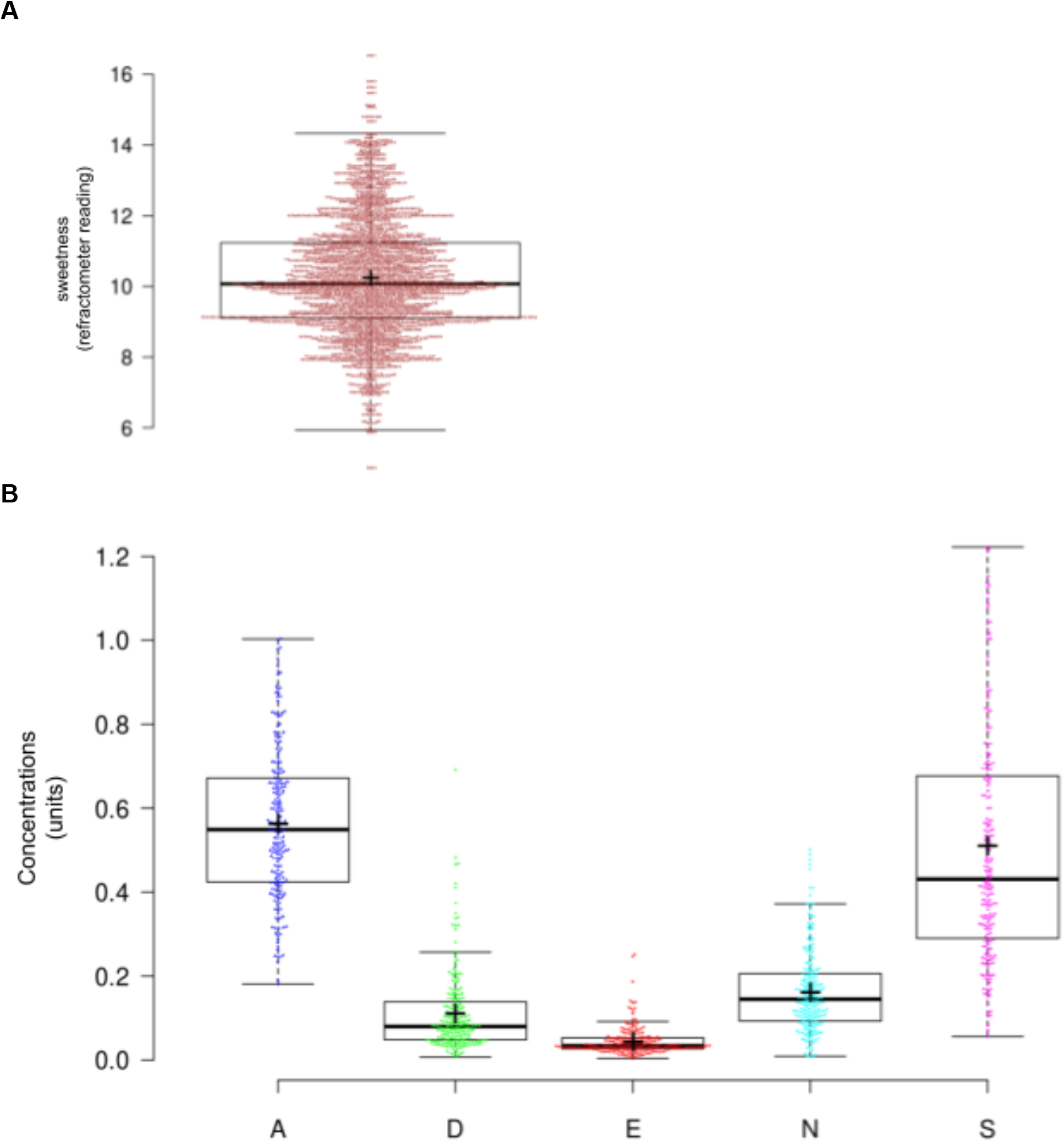
Distribution of measured analyte values. Refractometer readings of kinnows (A) and concentrations of A, D, E, N and S neem metabolites (B) are plotted as boxplots with the data points overlaid as a bee swarm scatter. Center lines show the medians; box limits indicate the 25th and 75th percentiles; whiskers extend 1.5 times the interquartile range from the 25th and 75th percentiles; outliers are represented by dots; crosses represent sample means; data points are plotted as open circles.

### Assessment of adequacy of training set by imputation

We found that down-sizing the data even by 10% has an immediate effect on the validation F1-score (**Figure 3**) and the other validation metrics (**Figure S1**). This did not change much across the additional down-sizing datasets up to 80%, beyond which at 90%, a noticeable effect could be observed where the validation F1 score dropped upto 0.66, accompanied with a concomitant decrease in the mAP metrics (**Figure 3** and **Figure S1**).

**Figure 3:**
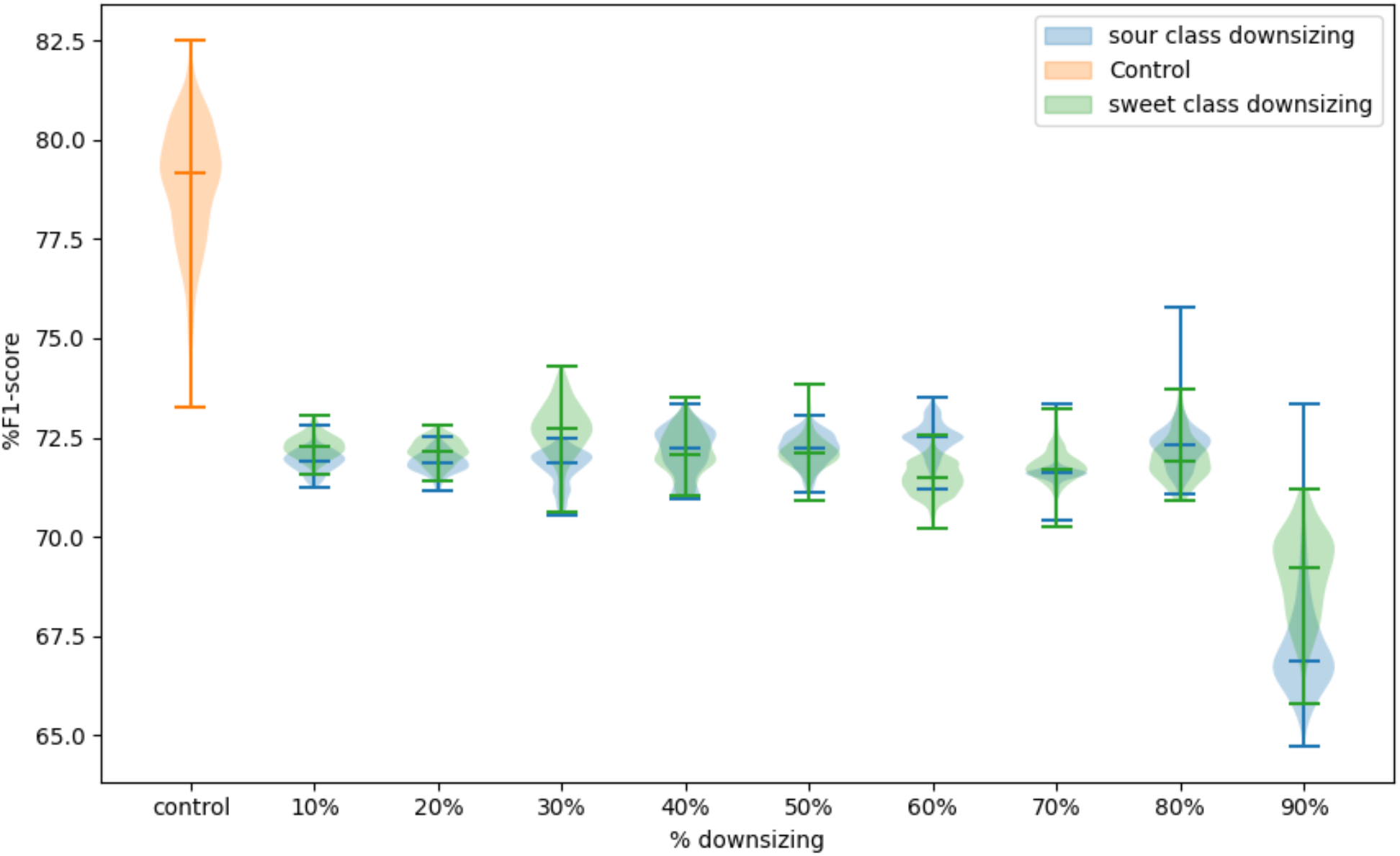
Distribution of validation F1-score in control versus serial class-wise down-sizing. Center lines in the violin plots show the medians; box limits indicate the 25th and 75th percentiles as determined by R software; whiskers extend 1.5 times the interquartile range from the 25th and 75th percentiles.

### Optimum training parameters

We observed F1-scores of 0.8 and 0.67 when training, validating and testing with 80:10:10 and 60:30:10 data splits for the kinnow model, and fixed the split to 80:10:10 for all our analyses. Further, we observed F1-scores of 0.663, 0.784 and 0.805 when training with batch sizes of 8, 32 and 64 on the kinnow images, and therefore, fixed the batch size to 64. Lastly, when the models were varied between small, medium and large categories, we observed similar F1-scores of 0.717, 0.719 and 0.718. Nevertheless, we fixed the model as the ‘medium’ variant for all analyses.

### Training and validation metrics

The metrics of performance for the kinnow ‘W model and neem ‘A’, ‘D’, ‘E’, ‘N’ and ‘S’ models before and after evolution of hyper-parameters are provided in **Table 2**. Most metrics except recall were the same or worse for training after tuning of hyper-parameters. This effect was more pronounced for four of the neem models, ‘A’, ‘D’, ‘E’ and ‘S’. Visuals of the metrics and losses for all the models are summarized in **Figure S2**. The variation in the class-independent and ‘high’ class-specific F1 scores among the ten randomly shuffled datasets created as part of the ten-fold cross-validation was less than 10% for the A, D and S models, while the variation in ‘low’ class-specific F1 scores across the ten randomly shuffled datasets was ~15% for all the 5 analyte models (**Figure 4**).

**Table 2:** Validation metrics of the best models for analyte classes before (A) and after (B) hyper-parameter tuning.

| *Fruit (analyte)* | *Class* | *Precision* | *Recall* | *mAP@.5* | *mAP@.5:.95* | *F1* |
| --- | --- | --- | --- | --- | --- | --- |
| *Kinnow sweetness (W)* | *All* | 0.72 | 0.88 | 0.86 | 0.83 | 0.79 |
|  | *Sweet* | 0.75 | 0.89 | 0.87 | 0.84 | 0.81 |
|  | *Sour* | 0.70 | 0.87 | 0.86 | 0.83 | 0.77 |
| *Neem (A)* | *All* | 0.60 | 0.89 | 0.76 | 0.70 | 0.72 |
|  | *High* | 0.68 | 0.82 | 0.78 | 0.72 | 0.75 |
|  | *Low* | 0.52 | 0.95 | 0.75 | 0.69 | 0.67 |
| *Neem (D)* | *All* | 0.62 | 0.92 | 0.80 | 0.74 | 0.74 |
|  | *High* | 0.58 | 0.92 | 0.82 | 0.77 | 0.71 |
|  | *Low* | 0.66 | 0.92 | 0.77 | 0.71 | 0.77 |
| *Neem (E)* | *All* | 0.61 | 0.91 | 0.79 | 0.74 | 0.73 |
|  | *High* | 0.47 | 0.93 | 0.75 | 0.70 | 0.63 |
|  | *Low* | 0.75 | 0.89 | 0.84 | 0.78 | 0.81 |
| *Neem (N)* | *All* | 0.71 | 0.73 | 0.74 | 0.70 | 0.72 |
|  | *High* | 0.88 | 0.55 | 0.82 | 0.77 | 0.68 |
|  | *Low* | 0.54 | 0.90 | 0.66 | 0.63 | 0.68 |
| *Neem (S)* | *All* | 0.75 | 0.86 | 0.89 | 0.81 | 0.80 |
|  | *High* | 0.71 | 0.94 | 0.87 | 0.81 | 0.81 |
|  | *Low* | 0.79 | 0.78 | 0.91 | 0.81 | 0.78 |

**B**
| *Fruit*<br>*(analyte)* | *Class* | *Precision* | *Recall* | *mAP@.5* | *mAP@.5:.95* | *F1* |
| --- | --- | --- | --- | --- | --- | --- |
| *Kinnow*<br>*sweetness*<br>*(W)* | - | 0.61 | 0.91 | 0.792 | 0.72 | 0.73 |
|  | *Sweet* | 0.64 | 0.92 | 0.80 | 0.73 | 0.75 |
|  | *Sour* | 0.59 | 0.9 | 0.79 | 0.72 | 0.71 |
| *Neem (A)* | - | 0.50 | 1.00 | 0.54 | 0.51 | 0.67 |
|  | *High* | 0.47 | 1.00 | 0.48 | 0.44 | 0.64 |
|  | *Low* | 0.53 | 1.00 | 0.60 | 0.58 | 0.69 |
| *Neem (D)* | - | 0.50 | 1.00 | 0.64 | 0.60 | 0.67 |
|  | *High* | 0.47 | 1.00 | 0.64 | 0.59 | 0.64 |
|  | *Low* | 0.53 | 1.00 | 0.64 | 0.61 | 0.69 |
| *Neem (E)* | - | 0.50 | 1.00 | 0.59 | 0.55 | 0.67 |
|  | *High* | 0.58 | 1.00 | 0.76 | 0.71 | 0.73 |
|  | *Low* | 0.42 | 1.00 | 0.42 | 0.40 | 0.59 |
| *Neem (N)* | - | 0.67 | 0.81 | 0.74 | 0.69 | 0.73 |
|  | *High* | 0.54 | 0.95 | 0.64 | 0.58 | 0.69 |
|  | *Low* | 0.80 | 0.66 | 0.85 | 0.79 | 0.73 |
| *Neem (S)* | - | 0.51 | 1.00 | 0.69 | 0.66 | 0.68 |
|  | *High* | 0.53 | 1.00 | 0.77 | 0.73 | 0.69 |
|  | *Low* | 0.49 | 1.00 | 0.61 | 0.59 | 0.66 |

**Figure 4:**
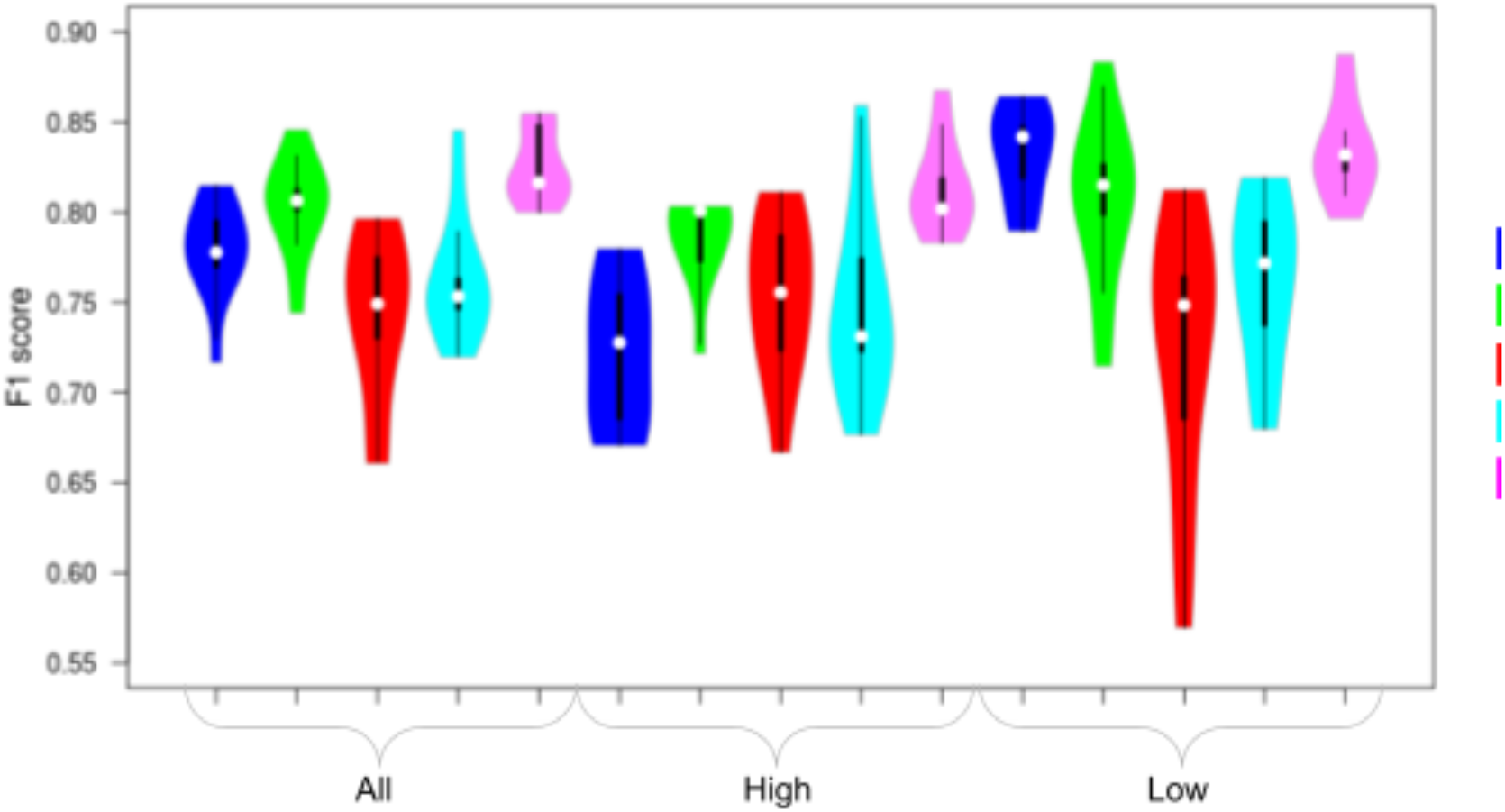
Distribution of 10-fold cross-validation F1 scores for all the neem models for all, high and low classes: White circles show the medians; box limits indicate the 25th and 75th percentiles; whiskers extend 1.5 times the interquartile range from the 25th and 75th percentiles; polygons represent density estimates of data and extend to extreme values.

### Test classification accuracy before and after boosting using the multi-analyte prediction model

We obtained test classification accuracy of 82.55% (82.18% for the sour class and 82.89% for the sweet class) for the kinnow model and 60.8% (64% for the low class and 56% for the high class) for the neem A model on the blind test set images. Further, the classification accuracy ranged from 60.8% to 80.2% (64% to 95% for the low class with a median value of 79.6% and 51.22% to 85.37% for the high class with a median value of 80.2%) across the ten shuffled cross validation datasets. The predictions on a few representative test set images for kinnow and neem fruit along with the confidence scores of the predictions are illustrated in **Figure 5** and on all images are provided in **Table S1** along with the actual classes. Here, in the case of the neem fruit, the same images result in five high or low class predictions using the ‘A’, ‘D’,’E’, ‘N’, ‘S’ models. Further, we used the advantage offered by these multi-analyte predictions for the same set of images to boost the prediction accuracy of the ‘A’ model. We identified a high:low:high prediction combination for D:N:S metabolites to map to an average of 18.6% and 2.93% of high and low ‘A’ class fruits, respectively, and a low:high prediction combination for D:N metabolites to map to an average of 5.33% and 31.11% of high and low ‘A’ class fruits, respectively. These combinations matched the high and low ‘A’ class fruits, respectively, with the highest sensitivity and specificity. When we replaced the A model predictions with the predictions as per these combinations, we observed a boosted classification accuracy ranging from 62.4% to 80.4% (67.8% to 95.1% for the low class with a median value of 81.4% and 48% to 83.33% for the high class with a median value of 60.5%) across the ten shuffles (**Figure 6**). The ten-fold cross-validation prediction error after the boosting was estimated at 0.257±0.057.

**Figure 5:**
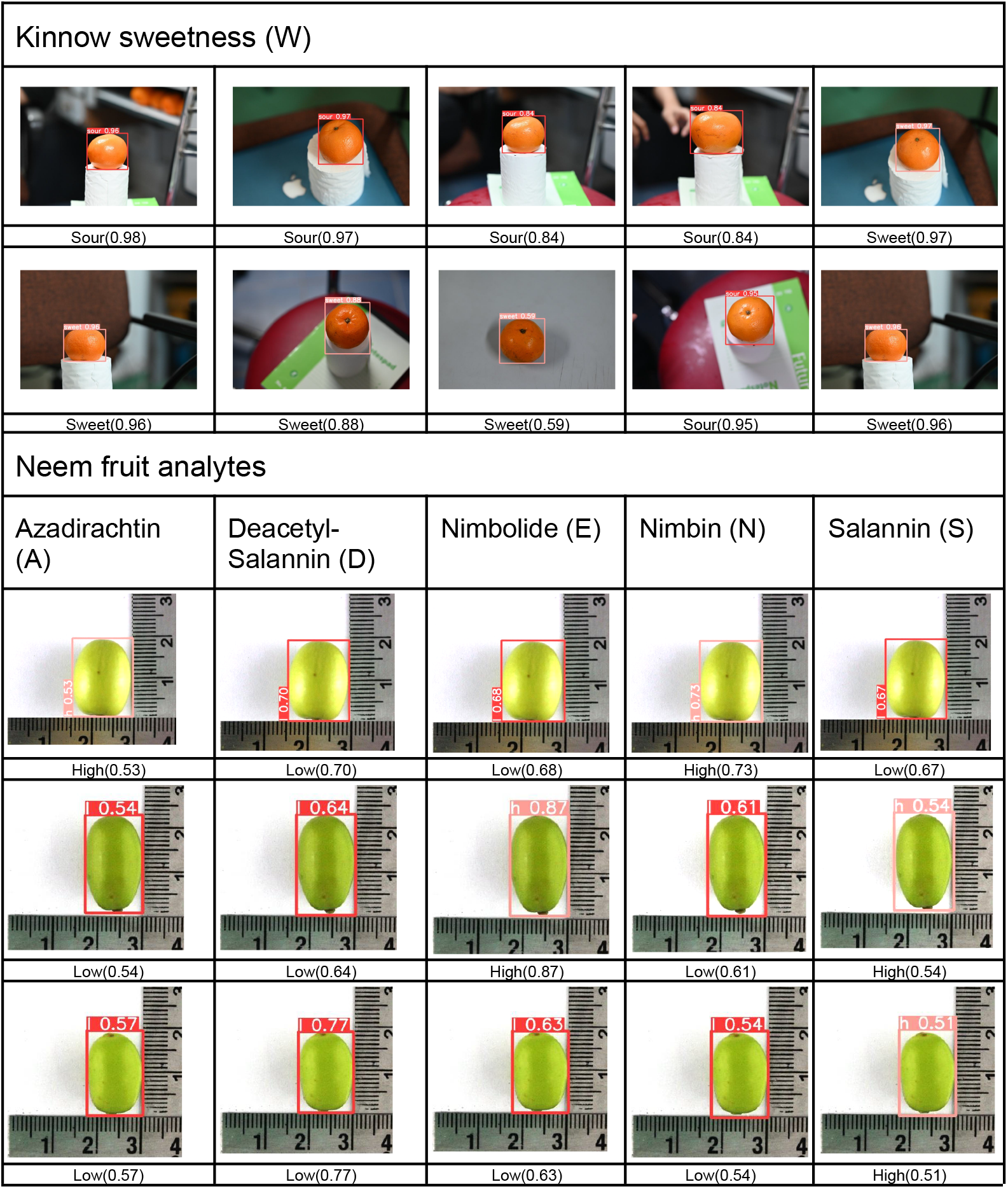
Prediction of fruit analyte classes in test set images. Representative images are shown here. The predicted bounding box and class along with the confidence score of prediction are displayed along with the corresponding image. Same images are shown horizontally across all neem fruit analyte models.

**Figure 6:**
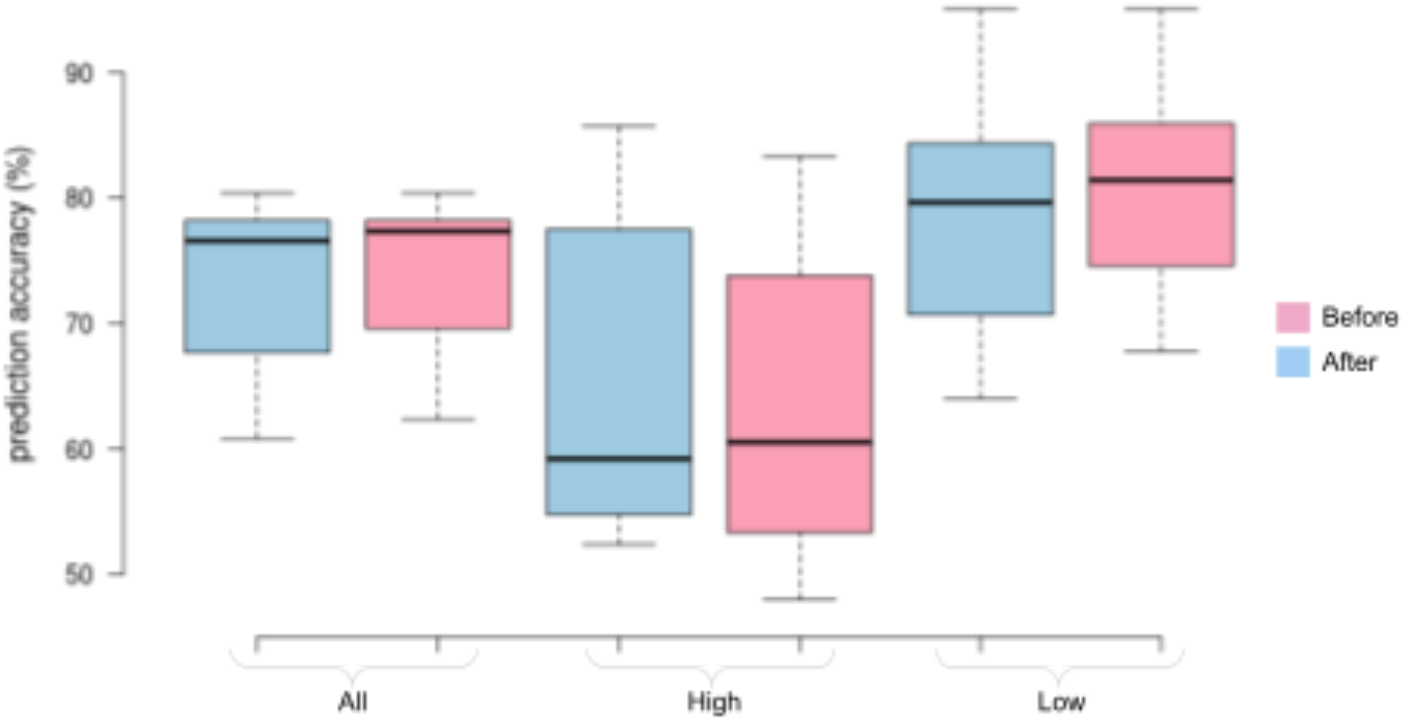
Neem ‘A’ model’s prediction accuracy (%) on test images for all, low and high classes: before and after multi-analyte boosting. The box plots represent the distribution of accuracies across ten random shuffles of the data. Center lines show the medians; box limits indicate the 25th and 75th percentiles; whiskers extend to minimum and maximum values.

## Discussion

Deep learning-based methods are producing impressive results across several domains, especially, visual and auditory recognition (LeCun et al. 2015). Data-intensive biological problems are well suited for deep learning methods (Ching et al. 2018). Biologically inspired neural networks are a class of machine learning algorithms that enable learning from data. Deep learning requires a neural network with multiple layers. Based on the nature of data and the type of question being asked, deep learning methods use either supervised, or unsupervised, or reinforcement learning-based training models. Convolutional neural networks (CNNs or ConvNets) are multi-layered neural networks trained with back-propagation algorithms, used in recognizing images with minimum pre-processing. In 1998, LeCun and co-workers had described a machine learning technique that was built to learn automatically with less hand-designed heuristics (LeCun et al. 1998). This formed the basis for development of the CNN field. CNNs combine three architectural ideas: local receptive fields, shared weights and, sometimes, spatial or temporal sub-sampling (LeCun et al. 1995). Biological data is often complex, multi-dimensional and heterogeneous. However, the possibility of using deep learning methods to discover patterns in such large and complex biological datasets is promising. Till date, image-based analysis have been used to study plant stress, phenotyping (Pound et al. 2017a; Pound et al. 2017b; Ghosal et al. 2018), and assess quality of fruits, grains, and vegetables (Al-Sammarraie et al. 2022; Boniecki et al. 2015; Zaborowicz et al. 2017; Koszela et al. 2015).

In our study, we used fruits from two trees, different in shape, size, color, and external texture, to image and used the images to test whether a deep learning-based approach can classify fruits into sweet and sour classes in one (kinnow) and predict high-metabolite bearing fruits (neem) in other. We measured the sugar content of kinnow using a refractometer. Sweetness as measured by soluble sugar is one of the parameters in fruit’s overall flavor and flavor is primarily attributed to the chemical composition of fruits such as sugars, acids and other chemical compounds. For secondary metabolites, we measured five of the important secondary metabolites from neem to provide endpoints for fruits’ agronomic and pharmaceutical importance.

The overall accuracy of our model for determining sweetness in kinnow fruits was 81% (**Table 1**). From the imputation experiments, we observed that downsizing the total data even by 10% (**Figure 3**), drastically reduced the metrics, in particular F1-score by 10% and mean absolute precision by 15%. Interestingly, downsizing beyond this even upto 80% had nearly the same impact as 10%. Only after downsizing data by 90% could a visible drop in metrics be observed by another 10-15%. Although these experiments could only be performed on the kinnow data, what it implies for the sparser neem fruit data where the image numbers are equivalent to the 60% downsizing, is that the F1-score is perhaps in the 10% reduced range. This conclusion assumes that the all fruit image classifications work identically, which is not the case. Previous studies attempted predicting flavor in fruits using linear and partial least square regression (Abegaz et al. 2004; Schwieterman et al. 2014; Gilbert et al. 2015), and random-forest (Eggink et al. 2012) with variable accuracies. In a recent study using tomato and blueberry (Colantonio et al. 2022), the authors have found that machine learning models using metabolome and sensory panel levels with multiple measurements from fruits can identify the optimum model that can predict consumer behavior. In their study, the authors have found that gradient-boosting machines and XGBoost models were most predictive (Colantonio et al. 2022). Another previous study using *k-*means clustering in watermelons yielded 84.62% accuracy (Nazulan et al. 2020). However, the study included only a limited number of fruits (*n* = 13). Al-Sammarraie and co-workers (Al-Sammarraie et al. 2022) compared various methods and found that the logistical regression analysis provided the highest accuracy (97%) to predict sweetness in oranges. In the study by Al-Sammarraie and co-workers, multiple methods provided an accuracy between 82-97%. We believe that the number of fruits in the above study (*n* = 50 overall and *n* = 8 for very high sweet class) were limiting for any deep learning-based method to correctly assess accuracy. Additionally, the authors pre-selected the fruits for sizes and color. In addition to single and multi-analyte models, we also tried regression-based deep learning models. Although preliminary, the initial results of using a regression model did not provide a high validation accuracy for kinnow classification, which might have resulted due to a high-degree of false positives (overfitting of the model). Unlike some of the previous studies as mentioned above, our study had a lot more fruit images (*n* = 3451) and the fruits were randomly selected from different markets representing diverse geographical regions of the trees, climate and growing conditions. In neem, the image classification yielded an accuracy of 75% using a single metabolite model, for azadirachtin (**Table 1**). In addition to azadirachtin, we also measured four other metabolites, sallinin, de-acetyl sallinin, nimbin and nimbolide, from the same fruits. When the model was trained with a single metabolite, it yielded a prediction accuracy between 63 - 81% for either of the two classes. As agronomic importance is attributed to multiple metabolites, we tried to boost the model’s accuracy by taking into account multiple analytes measured from the same fruits. Although not to a great extent, the overall class prediction accuracy was higher with the multi-analyte boosting (**Figure 6**). Future studies involving a multi-compound model may enhance the model’s predictive accuracy further. Additionally, like in kinnow, we tried a deep learning-based regression model with neem fruits where our preliminary findings did not yield satisfactory results. Currently, we are working to improve the accuracy of the regression-based prediction model further in neem fruits.

Despite the encouraging results, our study has several limitations. In kinnow, we used a crude soluble sugar measurement using a refractometer. It is most likely that there were multiple factors, beyond sugars, responsible for flavor in kinnow. Future research involving multiple factors, like acids and other volatile substances in kinnow, alongside sugar, may prove to be more accurate and predictive for the flavor. Second, in the case of neem, we achieved an accuracy of 75% with azadirachtin model alone and 79% when boosted with four other metabolites. The predictive power of deep learning-based methods depends on the number of images used for training. Although we had far more images (*n* = 3,451 for kinnow and *n* = 1,045 for neem) than some of the previous studies reported on various fruit sweetness models, the numbers are still low in the context of deep learning. For studies like ours, where images are linked with biological parameters, and unlike popular computer vision problems like facial recognition, it is time consuming, cumbersome and expensive to obtain a large number of images. This is especially true where a linked biological metabolite measurement is involved. In both kinnow and neem, as we did not perform any variance decomposition study, and therefore, it is likely that multiple factors, beyond sugar and a few metabolites, are linked with the visible trait. Like a recent study involving tomato and blueberry (Colantonio et al. 2022), it is possible that the kinnow and neem fruit images alone were not reflective of flavor and agronomic values. Future research with multiple measurements from the same fruits, including acids, metabolites and other compounds, may improve deep learning-based image classification.

