## Supplementary material for "Single and multi-analyte deep learning-based analysis framework for class prediction in biological images": Supplmentary Tables and Figures

### *Supplementary Tables*

***Table S1:*** *Image-wise analyte predictions for test set kinnow and neem fruit images.*

| | *Predicted* | | | | | *Actual* | | | | | | *Predicted* | *Actual* |
|---|---|---|---|---|---|---|---|---|---|---|---|---|---|
| ***Neem*** | ***A*** | ***D*** | ***E*** | ***N*** | ***S*** | ***A*** | ***D*** | ***E*** | ***N*** | ***S*** | ***Kinnow*** | ***W*** | ***W*** |
| fruit-N-T1 | High | High | High | High | High | High | High | Low | High | High | fruit-K-T1 | sour | sour |
| fruit-N-T2 | High | High | High | High | High | High | High | High | High | High | fruit-K-T2 | sour | sour |
| fruit-N-T3 | High | High | High | High | High | High | High | High | High | High | fruit-K-T3 | sour | sour |
| fruit-N-T4 | High | High | High | High | High | High | High | High | High | High | fruit-K-T4 | sour | sour |
| fruit-N-T5 | Low | High | High | High | High | High | High | High | High | High | fruit-K-T5 | sour | sour |
| fruit-N-T6 | Low | High | High | High | High | High | High | High | High | High | fruit-K-T6 | sour | sour |
| fruit-N-T7 | High | High | High | High | High | Low | High | High | High | High | fruit-K-T7 | sour | sour |
| fruit-N-T8 | High | High | High | High | High | Low | High | High | High | High | fruit-K-T8 | sour | sour |
| fruit-N-T9 | High | High | High | High | High | Low | High | High | High | High | fruit-K-T9 | sour | sour |
| fruit-N-T10 | High | High | High | High | High | Low | High | High | High | High | fruit-K-T10 | sour | sour |
| fruit-N-T11 | High | High | High | High | High | Low | High | High | High | High | fruit-K-T11 | sour | sour |
| fruit-N-T12 | Low | High | High | High | High | Low | Low | Low | Low | Low | fruit-K-T12 | sour | sour |
| fruit-N-T13 | High | High | High | Low | High | High | High | Low | High | High | fruit-K-T13 | sour | sour |
| fruit-N-T14 | High | High | High | Low | High | High | High | High | Low | High | fruit-K-T14 | sour | sour |
| fruit-N-T15 | Low | High | High | Low | High | High | Low | Low | Low | High | fruit-K-T15 | sour | sour |
| fruit-N-T16 | Low | High | High | Low | High | High | High | High | High | High | fruit-K-T16 | sour | sour |
| fruit-N-T17 | Low | High | High | Low | High | High | High | High | High | High | fruit-K-T17 | sweet | sour |
| fruit-N-T18 | Low | High | High | Low | High | High | High | High | Low | High | fruit-K-T18 | sour | sour |
| fruit-N-T19 | Low | High | High | Low | High | High | High | High | Low | High | fruit-K-T19 | sour | sour |
| fruit-N-T20 | Low | High | High | Low | High | High | High | High | Low | High | fruit-K-T20 | sour | sour |
| fruit-N-T21 | Low | High | High | Low | High | High | High | High | Low | High | fruit-K-T21 | sour | sour |
| fruit-N-T22 | Low | High | High | Low | High | Low | High | High | Low | High | fruit-K-T22 | sour | sour |
| fruit-N-T23 | High | High | High | Low | Low | High | Low | Low | Low | High | fruit-K-T23 | sour | sour |
| fruit-N-T24 | Low | High | Low | High | High | High | High | High | High | High | fruit-K-T24 | sour | sour |
| fruit-N-T25 | Low | High | Low | High | High | High | High | High | High | High | fruit-K-T25 | sour | sour |
| fruit-N-T26 | High | High | Low | Low | High | High | High | Low | High | High | fruit-K-T26 | sour | sour |
| fruit-N-T27 | Low | High | Low | Low | High | High | High | High | High | High | fruit-K-T27 | sweet | sour |
| fruit-N-T28 | Low | High | Low | Low | High | High | High | Low | High | High | fruit-K-T28 | sweet | sour |
| fruit-N-T29 | Low | High | Low | Low | High | High | High | Low | High | High | fruit-K-T29 | sour | sour |
| fruit-N-T30 | High | High | Low | Low | High | Low | High | High | Low | High | fruit-K-T30 | sour | sour |
| fruit-N-T31 | Low | High | Low | Low | High | Low | High | High | Low | High | fruit-K-T31 | sour | sour |
| fruit-N-T32 | Low | High | Low | Low | High | Low | High | High | Low | High | fruit-K-T32 | sour | sour |
| fruit-N-T33 | Low | High | Low | Low | High | Low | Low | Low | Low | Low | fruit-K-T33 | sour | sour |
| fruit-N-T34 | High | Low | High | High | Low | Low | Low | Low | Low | Low | fruit-K-T34 | sour | sour |
| fruit-N-T35 | High | Low | High | High | Low | Low | Low | Low | Low | Low | fruit-K-T35 | sour | sour |
| fruit-N-T36 | High | Low | High | High | Low | Low | Low | Low | Low | Low | fruit-K-T36 | sour | sour |
| fruit-N-T37 | High | Low | High | High | Low | Low | Low | Low | Low | Low | fruit-K-T37 | sour | sour |
| fruit-N-T38 | Low | Low | High | High | Low | Low | High | High | High | Low | fruit-K-T38 | sweet | sour |
| fruit-N-T39 | Low | Low | High | High | Low | Low | Low | High | Low | Low | fruit-K-T39 | sweet | sour |
| fruit-N-T40 | Low | Low | High | High | Low | Low | Low | High | Low | Low | fruit-K-T40 | sour | sour |
| fruit-N-T41 | Low | Low | High | High | Low | Low | Low | High | Low | Low | fruit-K-T41 | sour | sour |

| | | | | | | | | | | | | | |
|---|---|---|---|---|---|---|---|---|---|---|---|---|---|
| fruit-N-T42 | Low | Low | High | High | Low | Low | Low | High | Low | Low | fruit-K-T42 | sweet | sour |
| fruit-N-T43 | Low | Low | High | High | Low | Low | Low | High | Low | Low | fruit-K-T43 | sour | sour |
| fruit-N-T44 | Low | Low | High | Low | High | High | Low | Low | Low | High | fruit-K-T44 | sour | sour |
| fruit-N-T45 | High | Low | High | Low | High | Low | Low | Low | Low | Low | fruit-K-T45 | sour | sour |
| fruit-N-T46 | Low | Low | High | Low | High | Low | High | High | Low | High | fruit-K-T46 | sour | sour |
| fruit-N-T47 | Low | Low | High | Low | High | Low | Low | Low | Low | Low | fruit-K-T47 | sour | sour |
| fruit-N-T48 | High | Low | High | Low | Low | High | High | Low | High | High | fruit-K-T48 | sour | sour |
| fruit-N-T49 | Low | Low | High | Low | Low | Low | Low | Low | High | Low | fruit-K-T49 | sour | sour |
| fruit-N-T50 | Low | Low | High | Low | Low | Low | Low | Low | High | Low | fruit-K-T50 | sour | sour |
| fruit-N-T51 | Low | Low | High | Low | Low | Low | Low | High | Low | High | fruit-K-T51 | sour | sour |
| fruit-N-T52 | High | Low | Low | High | High | High | High | High | High | Low | fruit-K-T52 | sour | sour |
| fruit-N-T53 | High | Low | Low | High | High | Low | Low | Low | Low | Low | fruit-K-T53 | sour | sour |
| fruit-N-T54 | High | Low | Low | High | High | Low | Low | Low | Low | Low | fruit-K-T54 | sour | sour |
| fruit-N-T55 | High | Low | Low | High | High | Low | Low | Low | Low | Low | fruit-K-T55 | sour | sour |
| fruit-N-T56 | High | Low | Low | High | High | Low | High | High | High | Low | fruit-K-T56 | sour | sour |
| fruit-N-T57 | High | Low | Low | High | Low | High | High | High | Low | High | fruit-K-T57 | sour | sour |
| fruit-N-T58 | High | Low | Low | High | Low | High | High | High | High | Low | fruit-K-T58 | sour | sour |
| fruit-N-T59 | High | Low | Low | High | Low | High | High | High | High | Low | fruit-K-T59 | sour | sour |
| fruit-N-T60 | High | Low | Low | High | Low | High | High | High | High | Low | fruit-K-T60 | sour | sour |
| fruit-N-T61 | High | Low | Low | High | Low | High | High | High | High | Low | fruit-K-T61 | sweet | sour |
| fruit-N-T62 | High | Low | Low | High | Low | High | Low | High | High | Low | fruit-K-T62 | sour | sour |
| fruit-N-T63 | High | Low | Low | High | Low | High | Low | High | High | Low | fruit-K-T63 | sour | sour |
| fruit-N-T64 | High | Low | Low | High | Low | High | Low | High | High | Low | fruit-K-T64 | sour | sour |
| fruit-N-T65 | High | Low | Low | High | Low | High | Low | High | High | Low | fruit-K-T65 | sour | sour |
| fruit-N-T66 | High | Low | Low | High | Low | High | Low | High | High | Low | fruit-K-T66 | sweet | sour |
| fruit-N-T67 | High | Low | Low | High | Low | High | High | High | Low | Low | fruit-K-T67 | sour | sour |
| fruit-N-T68 | High | Low | Low | High | Low | High | High | High | Low | Low | fruit-K-T68 | sour | sour |
| fruit-N-T69 | High | Low | Low | High | Low | High | High | High | Low | Low | fruit-K-T69 | sour | sour |
| fruit-N-T70 | Low | Low | Low | High | Low | High | High | Low | High | High | fruit-K-T70 | sour | sour |
| fruit-N-T71 | Low | Low | Low | High | Low | High | High | High | Low | Low | fruit-K-T71 | sour | sour |
| fruit-N-T72 | Low | Low | Low | High | Low | High | High | High | Low | Low | fruit-K-T72 | sweet | sour |
| fruit-N-T73 | High | Low | Low | High | Low | Low | Low | Low | Low | Low | fruit-K-T73 | sour | sour |
| fruit-N-T74 | High | Low | Low | High | Low | Low | Low | Low | Low | Low | fruit-K-T74 | sour | sour |
| fruit-N-T75 | High | Low | Low | High | Low | Low | High | High | High | Low | fruit-K-T75 | sour | sour |
| fruit-N-T76 | High | Low | Low | High | Low | Low | Low | Low | Low | Low | fruit-K-T76 | sour | sour |
| fruit-N-T77 | High | Low | Low | High | Low | Low | Low | Low | Low | Low | fruit-K-T77 | sour | sour |
| fruit-N-T78 | High | Low | Low | High | Low | Low | Low | Low | Low | Low | fruit-K-T78 | sour | sour |
| fruit-N-T79 | High | Low | Low | High | Low | Low | Low | High | Low | High | fruit-K-T79 | sour | sour |
| fruit-N-T80 | High | Low | Low | High | Low | Low | Low | Low | Low | Low | fruit-K-T80 | sour | sour |
| fruit-N-T81 | High | Low | Low | High | Low | Low | Low | Low | Low | Low | fruit-K-T81 | sweet | sour |
| fruit-N-T82 | High | Low | Low | High | Low | Low | Low | Low | Low | Low | fruit-K-T82 | sweet | sour |
| fruit-N-T83 | High | Low | Low | High | Low | Low | Low | Low | Low | Low | fruit-K-T83 | sour | sour |
| fruit-N-T84 | Low | Low | Low | High | Low | Low | High | High | High | Low | fruit-K-T84 | sour | sour |
| fruit-N-T85 | Low | Low | Low | High | Low | Low | High | High | High | Low | fruit-K-T85 | sour | sour |
| fruit-N-T86 | Low | Low | Low | High | Low | Low | Low | High | Low | High | fruit-K-T86 | sweet | sour |
| fruit-N-T87 | Low | Low | Low | High | Low | Low | Low | High | Low | High | fruit-K-T87 | sour | sour |
| fruit-N-T88 | Low | Low | Low | High | Low | Low | Low | High | Low | High | fruit-K-T88 | sour | sour |
| fruit-N-T89 | Low | Low | Low | High | Low | Low | Low | High | Low | High | fruit-K-T89 | sour | sour |

| | | | | | | | | | | | | |
|---|---|---|---|---|---|---|---|---|---|---|---|---|
| fruit-N-T90 | Low | Low | Low | High | Low | Low | Low | High | Low | High | fruit-K-T90 | sour | sour |
| fruit-N-T91 | Low | Low | Low | High | Low | Low | Low | High | Low | High | fruit-K-T91 | sour | sour |
| fruit-N-T92 | Low | Low | Low | High | Low | Low | Low | High | Low | High | fruit-K-T92 | sour | sour |
| fruit-N-T93 | Low | Low | Low | High | Low | Low | Low | High | Low | High | fruit-K-T93 | sour | sour |
| fruit-N-T94 | Low | Low | Low | High | Low | Low | Low | High | Low | Low | fruit-K-T94 | sweet | sour |
| fruit-N-T95 | Low | Low | Low | High | Low | Low | Low | High | Low | Low | fruit-K-T95 | sour | sour |
| fruit-N-T96 | Low | Low | Low | High | Low | Low | Low | High | Low | Low | fruit-K-T96 | sour | sour |
| fruit-N-T97 | Low | Low | Low | High | Low | Low | Low | High | Low | Low | fruit-K-T97 | sour | sour |
| fruit-N-T98 | Low | Low | Low | High | Low | Low | Low | High | Low | Low | fruit-K-T98 | sweet | sour |
| fruit-N-T99 | Low | Low | Low | High | Low | Low | Low | Low | Low | Low | fruit-K-T99 | sweet | sour |
| fruit-N-T100 | Low | Low | Low | High | Low | Low | Low | Low | Low | Low | fruit-K-T100 | sour | sour |
| fruit-N-T101 | Low | Low | Low | High | Low | Low | Low | Low | Low | Low | fruit-K-T101 | sour | sour |
| fruit-N-T102 | Low | Low | Low | High | Low | Low | Low | Low | Low | Low | fruit-K-T102 | sour | sour |
| fruit-N-T103 | Low | Low | Low | High | Low | Low | Low | High | Low | Low | fruit-K-T103 | sour | sour |
| fruit-N-T104 | Low | Low | Low | High | Low | Low | Low | High | Low | Low | fruit-K-T104 | sour | sour |
| fruit-N-T105 | Low | Low | Low | High | Low | Low | Low | High | Low | Low | fruit-K-T105 | sour | sour |
| fruit-N-T106 | Low | Low | Low | High | Low | Low | Low | High | Low | Low | fruit-K-T106 | sour | sour |
| fruit-N-T107 | Low | Low | Low | High | Low | Low | Low | High | Low | Low | fruit-K-T107 | sour | sour |
| fruit-N-T108 | High | Low | Low | Low | High | High | Low | Low | Low | High | fruit-K-T108 | sour | sour |
| fruit-N-T109 | Low | Low | Low | Low | High | High | Low | Low | Low | High | fruit-K-T109 | sweet | sour |
| fruit-N-T110 | Low | Low | Low | Low | High | High | High | Low | High | High | fruit-K-T110 | sour | sour |
| fruit-N-T111 | High | Low | Low | Low | High | Low | Low | Low | Low | Low | fruit-K-T111 | sweet | sour |
| fruit-N-T112 | Low | Low | Low | Low | High | Low | Low | Low | Low | Low | fruit-K-T112 | sour | sour |
| fruit-N-T113 | Low | Low | Low | Low | High | Low | Low | Low | Low | Low | fruit-K-T113 | sour | sour |
| fruit-N-T114 | Low | Low | Low | Low | High | Low | Low | Low | Low | Low | fruit-K-T114 | sour | sour |
| fruit-N-T115 | Low | Low | Low | Low | High | Low | Low | Low | Low | Low | fruit-K-T115 | sour | sour |
| fruit-N-T116 | Low | Low | Low | Low | High | Low | Low | Low | Low | Low | fruit-K-T116 | sour | sour |
| fruit-N-T117 | High | Low | Low | Low | Low | High | High | High | Low | High | fruit-K-T117 | sweet | sour |
| fruit-N-T118 | High | Low | Low | Low | Low | High | High | High | Low | High | fruit-K-T118 | sour | sour |
| fruit-N-T119 | High | Low | Low | Low | Low | High | High | High | Low | High | fruit-K-T119 | sour | sour |
| fruit-N-T120 | High | Low | Low | Low | Low | High | High | High | Low | High | fruit-K-T120 | sweet | sour |
| fruit-N-T121 | Low | Low | Low | Low | Low | High | High | Low | High | High | fruit-K-T121 | sour | sour |
| fruit-N-T122 | Low | Low | Low | Low | Low | High | High | Low | High | High | fruit-K-T122 | sweet | sour |
| fruit-N-T123 | Low | Low | Low | Low | Low | Low | Low | Low | High | Low | fruit-K-T123 | sour | sour |
| fruit-N-T124 | Low | Low | Low | Low | Low | Low | Low | Low | High | Low | fruit-K-T124 | sweet | sour |
| fruit-N-T125 | Low | Low | Low | Low | Low | Low | Low | Low | High | Low | fruit-K-T125 | sour | sour |
| | | | | | | | | | | | fruit-K-T126 | sweet | sour |
| | | | | | | | | | | | fruit-K-T127 | sweet | sour |
| | | | | | | | | | | | fruit-K-T128 | sour | sour |
| | | | | | | | | | | | fruit-K-T129 | sour | sour |
| | | | | | | | | | | | fruit-K-T130 | sweet | sour |
| | | | | | | | | | | | fruit-K-T131 | sweet | sour |
| | | | | | | | | | | | fruit-K-T132 | sour | sour |
| | | | | | | | | | | | fruit-K-T133 | sour | sour |
| | | | | | | | | | | | fruit-K-T134 | sour | sour |
| | | | | | | | | | | | fruit-K-T135 | sour | sour |
| | | | | | | | | | | | fruit-K-T136 | sweet | sour |
| | | | | | | | | | | | fruit-K-T137 | sweet | sour |

| | | |
|---|---|---|
| fruit-K-T138 | sour | sour |
| fruit-K-T139 | sour | sour |
| fruit-K-T140 | sour | sour |
| fruit-K-T141 | sour | sour |
| fruit-K-T142 | sweet | sour |
| fruit-K-T143 | sour | sour |
| fruit-K-T144 | sour | sour |
| fruit-K-T145 | sweet | sour |
| fruit-K-T146 | sweet | sour |
| fruit-K-T147 | sour | sour |
| fruit-K-T148 | sour | sour |
| fruit-K-T149 | sour | sour |
| fruit-K-T150 | sour | sour |
| fruit-K-T151 | sweet | sour |
| fruit-K-T152 | sour | sour |
| fruit-K-T153 | sweet | sour |
| fruit-K-T154 | sour | sweet |
| fruit-K-T155 | sweet | sweet |
| fruit-K-T156 | sweet | sweet |
| fruit-K-T157 | sweet | sweet |
| fruit-K-T158 | sweet | sweet |
| fruit-K-T159 | sweet | sweet |
| fruit-K-T160 | sweet | sweet |
| fruit-K-T161 | sweet | sweet |
| fruit-K-T162 | sour | sweet |
| fruit-K-T163 | sour | sweet |
| fruit-K-T164 | sweet | sweet |
| fruit-K-T165 | sweet | sweet |
| fruit-K-T166 | sweet | sweet |
| fruit-K-T167 | sweet | sweet |
| fruit-K-T168 | sour | sweet |
| fruit-K-T169 | sour | sweet |
| fruit-K-T170 | sweet | sweet |
| fruit-K-T171 | sweet | sweet |
| fruit-K-T172 | sour | sweet |
| fruit-K-T173 | sweet | sweet |
| fruit-K-T174 | sweet | sweet |
| fruit-K-T175 | sweet | sweet |
| fruit-K-T176 | sweet | sweet |
| fruit-K-T177 | sweet | sweet |
| fruit-K-T178 | sweet | sweet |
| fruit-K-T179 | sour | sweet |
| fruit-K-T180 | sweet | sweet |
| fruit-K-T181 | sweet | sweet |
| fruit-K-T182 | sweet | sweet |
| fruit-K-T183 | sweet | sweet |
| fruit-K-T184 | sour | sweet |
| fruit-K-T185 | sour | sweet |

| | | |
|---|---|---|
| fruit-K-T186 | sweet | sweet |
| fruit-K-T187 | sweet | sweet |
| fruit-K-T188 | sweet | sweet |
| fruit-K-T189 | sweet | sweet |
| fruit-K-T190 | sour | sweet |
| fruit-K-T191 | sour | sweet |
| fruit-K-T192 | sweet | sweet |
| fruit-K-T193 | sweet | sweet |
| fruit-K-T194 | sweet | sweet |
| fruit-K-T195 | sweet | sweet |
| fruit-K-T196 | sweet | sweet |
| fruit-K-T197 | sweet | sweet |
| fruit-K-T198 | sweet | sweet |
| fruit-K-T199 | sweet | sweet |
| fruit-K-T200 | sour | sweet |
| fruit-K-T201 | sour | sweet |
| fruit-K-T202 | sweet | sweet |
| fruit-K-T203 | sour | sweet |
| fruit-K-T204 | sour | sweet |
| fruit-K-T205 | sweet | sweet |
| fruit-K-T206 | sweet | sweet |
| fruit-K-T207 | sweet | sweet |
| fruit-K-T208 | sweet | sweet |
| fruit-K-T209 | sweet | sweet |
| fruit-K-T210 | sweet | sweet |
| fruit-K-T211 | sweet | sweet |
| fruit-K-T212 | sour | sweet |
| fruit-K-T213 | sour | sweet |
| fruit-K-T214 | sweet | sweet |
| fruit-K-T215 | sweet | sweet |
| fruit-K-T216 | sweet | sweet |
| fruit-K-T217 | sour | sweet |
| fruit-K-T218 | sweet | sweet |
| fruit-K-T219 | sweet | sweet |
| fruit-K-T220 | sweet | sweet |
| fruit-K-T221 | sweet | sweet |
| fruit-K-T222 | sweet | sweet |
| fruit-K-T223 | sweet | sweet |
| fruit-K-T224 | sweet | sweet |
| fruit-K-T225 | sweet | sweet |
| fruit-K-T226 | sweet | sweet |
| fruit-K-T227 | sweet | sweet |
| fruit-K-T228 | sweet | sweet |
| fruit-K-T229 | sour | sweet |
| fruit-K-T230 | sweet | sweet |
| fruit-K-T231 | sweet | sweet |
| fruit-K-T232 | sweet | sweet |
| fruit-K-T233 | sour | sweet |

| | | |
|---|---|---|
| fruit-K-T234 | sweet | sweet |
| fruit-K-T235 | sweet | sweet |
| fruit-K-T236 | sweet | sweet |
| fruit-K-T237 | sour | sweet |
| fruit-K-T238 | sweet | sweet |
| fruit-K-T239 | sweet | sweet |
| fruit-K-T240 | sweet | sweet |
| fruit-K-T241 | sweet | sweet |
| fruit-K-T242 | sweet | sweet |
| fruit-K-T243 | sweet | sweet |
| fruit-K-T244 | sweet | sweet |
| fruit-K-T245 | sweet | sweet |
| fruit-K-T246 | sweet | sweet |
| fruit-K-T247 | sweet | sweet |
| fruit-K-T248 | sweet | sweet |
| fruit-K-T249 | sweet | sweet |
| fruit-K-T250 | sour | sweet |
| fruit-K-T251 | sour | sweet |
| fruit-K-T252 | sweet | sweet |
| fruit-K-T253 | sweet | sweet |
| fruit-K-T254 | sour | sweet |
| fruit-K-T255 | sweet | sweet |
| fruit-K-T256 | sweet | sweet |
| fruit-K-T257 | sweet | sweet |
| fruit-K-T258 | sour | sweet |
| fruit-K-T259 | sweet | sweet |
| fruit-K-T260 | sweet | sweet |
| fruit-K-T261 | sweet | sweet |
| fruit-K-T262 | sweet | sweet |
| fruit-K-T263 | sweet | sweet |
| fruit-K-T264 | sweet | sweet |
| fruit-K-T265 | sweet | sweet |
| fruit-K-T266 | sweet | sweet |
| fruit-K-T267 | sweet | sweet |
| fruit-K-T268 | sour | sweet |
| fruit-K-T269 | sour | sweet |
| fruit-K-T270 | sweet | sweet |
| fruit-K-T271 | sweet | sweet |
| fruit-K-T272 | sweet | sweet |
| fruit-K-T273 | sweet | sweet |
| fruit-K-T274 | sweet | sweet |
| fruit-K-T275 | sour | sweet |
| fruit-K-T276 | sweet | sweet |
| fruit-K-T277 | sweet | sweet |
| fruit-K-T278 | sweet | sweet |
| fruit-K-T279 | sweet | sweet |
| fruit-K-T280 | sweet | sweet |
| fruit-K-T281 | sweet | sweet |

| | | |
|---|---|---|
| fruit-K-T282 | sweet | sweet |
| fruit-K-T283 | sweet | sweet |
| fruit-K-T284 | sweet | sweet |
| fruit-K-T285 | sweet | sweet |
| fruit-K-T286 | sweet | sweet |
| fruit-K-T287 | sweet | sweet |
| fruit-K-T288 | sweet | sweet |
| fruit-K-T289 | sour | sweet |
| fruit-K-T290 | sweet | sweet |
| fruit-K-T291 | sweet | sweet |
| fruit-K-T292 | sweet | sweet |
| fruit-K-T293 | sweet | sweet |
| fruit-K-T294 | sweet | sweet |
| fruit-K-T295 | sweet | sweet |
| fruit-K-T296 | sweet | sweet |
| fruit-K-T297 | sweet | sweet |
| fruit-K-T298 | sweet | sweet |
| fruit-K-T299 | sweet | sweet |
| fruit-K-T300 | sweet | sweet |
| fruit-K-T301 | sweet | sweet |
| fruit-K-T302 | sweet | sweet |
| fruit-K-T303 | sweet | sweet |
| fruit-K-T304 | sweet | sweet |
| fruit-K-T305 | sweet | sweet |
| fruit-K-T306 | sweet | sweet |
| fruit-K-T307 | sweet | sweet |
| fruit-K-T308 | sweet | sweet |
| fruit-K-T309 | sweet | sweet |
| fruit-K-T310 | sweet | sweet |
| fruit-K-T311 | sour | sweet |
| fruit-K-T312 | sweet | sweet |
| fruit-K-T313 | sweet | sweet |
| fruit-K-T314 | sweet | sweet |
| fruit-K-T315 | sweet | sweet |
| fruit-K-T316 | sweet | sweet |
| fruit-K-T317 | sweet | sweet |
| fruit-K-T318 | sweet | sweet |
| fruit-K-T319 | sweet | sweet |
| fruit-K-T320 | sweet | sweet |
| fruit-K-T321 | sweet | sweet |
| fruit-K-T322 | sweet | sweet |
| fruit-K-T323 | sweet | sweet |
| fruit-K-T324 | sour | sweet |
| fruit-K-T325 | sweet | sweet |
| fruit-K-T326 | sweet | sweet |
| fruit-K-T327 | sweet | sweet |
| fruit-K-T328 | sweet | sweet |
| fruit-K-T329 | sweet | sweet |

| | | |
|---|---|---|
| fruit-K-T330 | sweet | sweet |
| fruit-K-T331 | sweet | sweet |
| fruit-K-T332 | sweet | sweet |
| fruit-K-T333 | sweet | sweet |
| fruit-K-T334 | sweet | sweet |
| fruit-K-T335 | sweet | sweet |
| fruit-K-T336 | sweet | sweet |
| fruit-K-T337 | sweet | sweet |
| fruit-K-T338 | sour | sour |
| fruit-K-T339 | sweet | sweet |
| fruit-K-T340 | sweet | sweet |
| fruit-K-T341 | sour | sour |
| fruit-K-T342 | sour | sour |
| fruit-K-T343 | sour | sour |
| fruit-K-T344 | sour | sour |
| fruit-K-T345 | sour | sour |
| fruit-K-T346 | sour | sour |
| fruit-K-T347 | sour | sour |
| fruit-K-T348 | sour | sour |
| fruit-K-T349 | sour | sour |
| fruit-K-T350 | sour | sour |
| fruit-K-T351 | sour | sour |
| fruit-K-T352 | sour | sour |
| fruit-K-T353 | sour | sour |
| fruit-K-T354 | sour | sour |
| fruit-K-T355 | sour | sour |
| fruit-K-T356 | sour | sour |
| fruit-K-T357 | sour | sour |
| fruit-K-T358 | sour | sour |
| fruit-K-T359 | sour | sour |
| fruit-K-T360 | sour | sour |
| fruit-K-T361 | sour | sour |

### ***Supplementary Figures***

***Figure S1:*** *Distribution of validation metrics. Precision (A), Recall (B), mAP:0.5 (C) and mAP:0.5:0.95 (D) in control versus serial class-wise down-sizing. Center lines in violin plots show the medians; box limits indicate the 25th and 75th percentiles; whiskers extend 1.5 times the interquartile range from the 25th and 75th percentiles.*

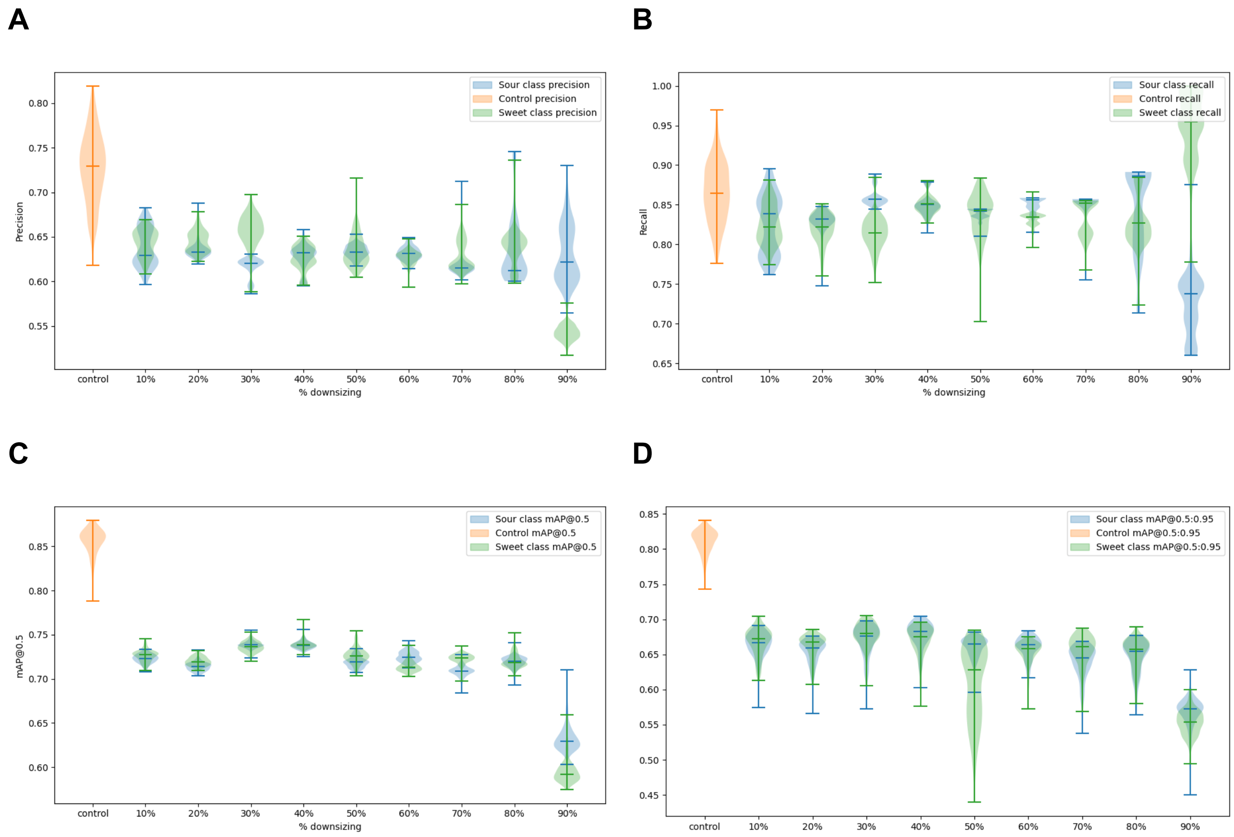

***Figure S2:*** *Performance metrics and loss curves for training and validation data for analyte classes in kinnow (A) and neem (B) fruits. The training and validation logs were sourced from Weights and Biases.*

A

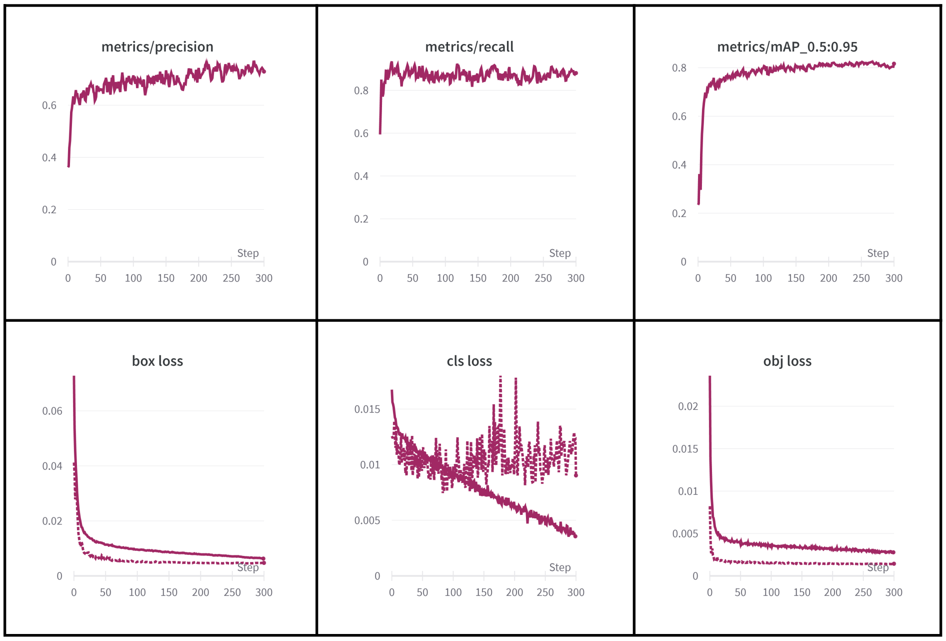


B

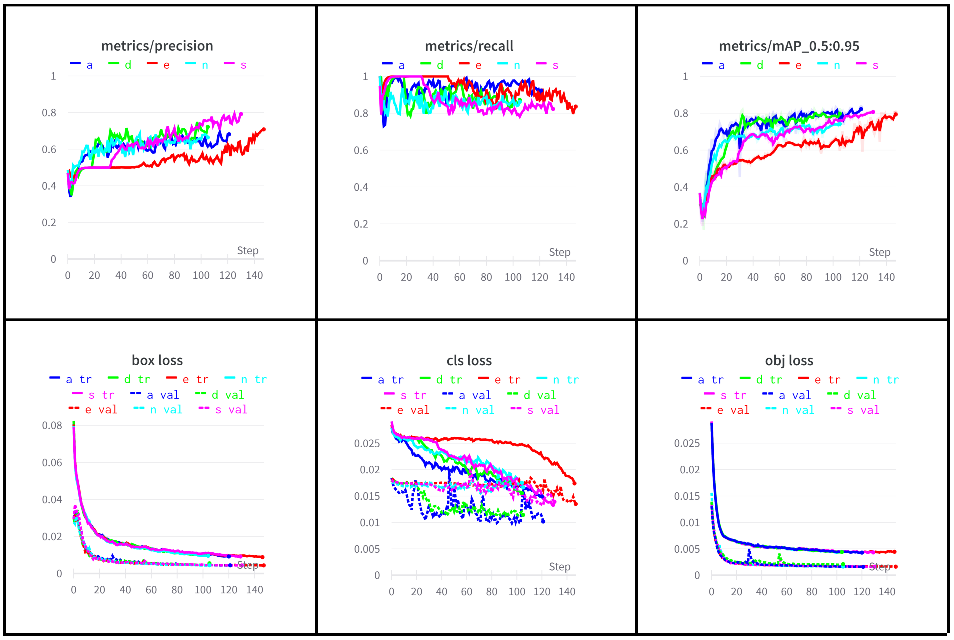
